# Gene Expression Detection in Developing Mouse Tissue Using In Situ Hybridization and µCT Imaging

**DOI:** 10.1101/2022.11.16.515587

**Authors:** Vilma Väänänen, Mona M. Christensen, Heikki Suhonen, Jukka Jernvall

## Abstract

High resolution and noninvasiveness have made soft tissue X-ray microtomography (µCT) a widely applicable three-dimensional (3D) imaging method in studies of morphology and development. However, scarcity of molecular probes to visualize gene activity with µCT has remained a challenge. Here we apply horseradish peroxidase -assisted reduction of silver and catalytic gold enhancement of the silver deposit to *in situ* hybridization in order to detect gene expression in developing tissues with µCT (here called GECT, Gene Expression CT). We show that GECT detects expression patterns of *collagen type II alpha 1 (Col2a1*) and *sonic hedgehog (Shh*) in developing mouse tissues comparably with an alkaline phosphatase-based detection method. After detection, expression patterns are visualized with laboratory µCT, demonstrating that GECT is compatible with varying levels of gene expression and varying sizes of expression regions. Additionally, we show that the method is compatible with prior phosphotungstic acid (PTA) staining, a conventional contrast staining approach in µCT imaging of soft tissues. Overall, GECT is a method that can be integrated with existing laboratory routines to obtain spatially accurate 3D detection of gene expression.

## Introduction

Studies of animal morphology and development require accurate visual information of three-dimensional structures that are typically too opaque for visible-light imaging. X-ray imaging methods are highly suitable for this purpose due to their penetrance and high, uniform resolution. One of the currently most accessible formats of 3D X-ray imaging is X-ray microtomography (µCT), which allows microscale resolution imaging of intact samples with relatively straightforward sample preparation using benchtop imaging equipment.

In X-ray attenuation µCT, X-ray projections of the sample are collected in various angles and then used to computationally generate a 3D model of how the sample reduces intensity of the X-rays traveling through it (Mizutani & Suzuki, 2012; Rawson et al., 2020). As the X-ray attenuation is considered proportional to the electron density and atomic number (Z) of the elements in imaged tissue, quantifiable volumetric data is obtained (Mizutani & Suzuki, 2012). 3D X-ray imaging techniques are readily applicable to nanoscale resolution imaging with techniques utilizing high-brightness synchrotron radiation, but nanoCT approaches without the need for synchrotron access are also currently available (Rawson et al., 2020). High-resolution X-ray imaging techniques allow phase contrast imaging, which has the practical benefit of samples not needing to go through any contrast enhancement procedures prior imaging (Mizutani & Suzuki, 2012; Rawson et al., 2020)

Due to intrinsic low X-ray attenuation of soft tissue, attenuation-based µCT using polychromatic X-ray sources often require procedures to increase contrast (Mizutani & Suzuki, 2012). This can be in some instances achieved by critical point drying (Zysk et al., 2012), but more widely utilized method is the use of high atomic number contrast agents (Mizutani & Suzuki, 2012; Pauwels et al., 2013)

Presence of the contrast agent in the tissue increases X-ray absorption of the sample in an energy-dependent manner (Mizutani & Suzuki, 2012; Rawson et al., 2020). Imaging a stained sample with X-ray energies near the k-shell electron binding energy of contrast agent atoms results in sudden increase in X-ray absorption, a phenomenon referred to as K-absorption edge (K-edge). K-edge increases as the atomic number of the element increases, and many of the established X-ray imaging applications rely on contrast agents with high atomic number. Commonly used contrast agents include iodine (Z=53, K-edge 33.2 keV) and tungsten (phosphotungstic acid, PTA, Z=74, K-edge 69.5 keV), both recommended due to their ease of use and low toxicity (Metscher, 2009b, 2009a; Pauwels et al., 2013). More recently, eosin, a common compound used in histology, was shown to generate high contrast suitable for high-resolution X-ray imaging of mouse kidney (Busse et al., 2018). Bromine (Z=35, K-edge 13.5 keV) present in eosin molecules generates X-ray attenuation during µCT imaging (Busse et al., 2018). Applying similar idea of utilizing conventional histological stains as µCT contrast agents, a method was described based on the use of hematein in complex with led (Z =82, K-edge 88 keV) for high-resolution nanoCT imaging of mouse liver and subsequent histological sectioning (Müller et al., 2018). Yet another contrast agent is protargol, a silver-protein compound that has been used as a general tissue contrast stain in synchrotron µCT imaging of tooth development (Raj et al., 2014). Silver has atomic number of 47 and K-edge of 25.5 keV, which translates to higher X-ray absorption in silver containing regions compared to unstained tissue when imaged with X-ray energies near 25.5 keV. Compared to conventional PTA staining, protargol enabled high quality imaging of early stage embryos, but suffered the same penetration problems as does PTA when imaging was extended to older whole embryo heads and intact embryos (Raj et al., 2014).

In contrast to the imaging of tissues, a long-standing challenge in biological µCT imaging has been the limited number of methods available to detect specific molecules or gene activity. Quite recently, enhancement of metal 3,3-diaminobenzidine detection (nickel/cobalt DAB) with osmium tetroxide and uranyl acetate was used in the visualization of Ret antibody in wild type and mutant mice (Hoshi et al., 2018). Enhanced DAB detection was also the basis of a method where use of peroxidase reporter gene enabled staining of specific cell populations in Drosophila legs and mouse brain (Kuan et al., 2020). Also, a LacZ reporter gene system was recently established for µCT imaging of gene expression, utilizing the radiodensity of bromine in the product of the β-galactosidase reaction (Ermakova et al., 2021).

Silver has demonstrated potential also as a more specific contrast stain for X-ray imaging. Recently, modified Fontana-Masson staining was used for 3D detection and quantification of melanin content in zebrafish larvae (Katz et al., 2021). Immunohistochemical approach by Metscher & Müller (2011) utilized peroxidase-facilitated reduction of silver ions on sites of target proteins, and the 3D distribution of collagen type 2 and acetylated alpha tubulin in chick embryos was then visualized with µCT imaging.

Despite these advances, detection of gene expression, a central focus of developmental biology studies, remains challenging using µCT in a way that can be integrated with existing laboratory routines. To this end, here were introduce a method for the detection of gene expression in messenger RNA (mRNA) level, utilizing peroxidase-assisted reduction of silver in *in situ* hybridization.

### A Gene Expression CT (GECT) method to detect gene expression in developing tissues

To explore the usefulness of silver-based methods for visualization of gene expression using attenuation µCT, we detected the expression of two genes, *collagen type 2 alpha 1* (*Col2a1*) and *sonic hedgehog (Shh)*, in embryonic tissues of mouse using silver reducing *in situ* hybridization. We then compared the resulting expression patterns to conventional chromogenic *in situ* hybridization done in parallel with silver detection (Fig 1A, B.)

In both *in situ* hybridization approaches, mRNA of interest is detected with digoxigenin (DIG)-labeled RNA-probes. Whereas in chromogenic *in situ* hybridization the detection is based on alkaline phosphatase activity, silver *in situ* hybridization utilizes anti-DIG antibody conjugated to horseradish peroxidase (HRP) (Fig 1C). HRP facilitates specific reduction of silver ions in solution on sites of gene activity. In the presence of reducer, elemental silver condensates are formed (Powell et al., 2002, 2007). (Fig 1C).

Compared to proteins distributed abundantly, the general challenge in imaging gene expression is the detection of transient and intracellular messenger RNA (mRNA). Silver has K-absorption edge in the lower end of the standard µCT energy spectrum, a challenge in case the molecule of interest is present in very low numbers. Therefore, to increase the X-ray absorption of the expression sites, we utilized a commercial gold enhancement kit (Nanoprobes) to strengthen the expression signal and compared the resulting expression patterns to the conventional chromogenic method (Fig 1D).

Gold enhancement is based on autometallography, where metal ions are catalytically reduced on the surface of already existing metal particles in the presence of a reducer (Danscher, 1984; Grizzle, 1996). Commercial gold enhancement protocols do not state exact components of the detection reaction (Hainfeld et al., 1999; Weipoltshammer & Schöfer, 2000), but contain a set of solutions called “enhancer”, “activator” and “initiator” and buffer (Hainfeld et al., 1999; Weipoltshammer & Schöfer, 2000). Sodium thiosulphate washes reduce the background staining and stop the detection reaction (Hainfeld et al., 1999). We aimed to enhance the silver precipitate generated by peroxidase assisted silver reduction with elemental gold (Z=79, K-edge 80.7 keV), and thus increase the electron density of the expression sites towards a range suitable for routine µCT visualization.

To test whether our approach can detect widespread expression fairly deep inside tissue, we conducted *in situ* hybridization for *collagen type 2 alpha 1* (*Col2a1*) for whole embryonic day 12.5 mouse embryo (Fig 1). This should provide largely comparable patterns to the type II collagen protein detected in embryonic chicken previously using µCT (Metscher & Müller, 2011). During later embryogenesis after embryonic day 12*, Col2a1* is a prominent marker of differentiating chondrocytes in the developing skeletal system (Cheah et al., 1991; Ng et al., 1997; Sandberg & Vuorio, 1987). More specifically, *Col2a1* is highly expressed in developing digits and limbs, developing vertebrae and ribs and developing parietal and occipital bones of skull (Sakai et al., 2001).

To challenge the method in conditions where the target molecule is expressed locally and in low numbers, we show that GECT can be used to visualize expression of *sonic hedgehog* (*Shh*) in a developing tooth germ of embryonic mouse (Fig 1). *Shh* is a central gene in the development of several mammalian organ systems, including ectodermal appendages such as teeth, hair buds and taste buds (Bitgood & McMahon, 1995; Dassule et al., 2000; Hallikas et al., 2021). During mouse embryonic day 14 to 15, *Shh* is expressed in the embryonic signaling center of a tooth germ, called the primary enamel knot (Jernvall et al., 1998; Jernvall et al., 1994).

To further establish the method as a way to visualize 3D gene expression and tissue morphology from an individual tissue sample, we combine phosphotungstic acid staining and µCT imaging of 3D morphology to the detection pipeline, from now on called GECT (Gene Expression CT) (Fig 1E).

**Figure 1.**
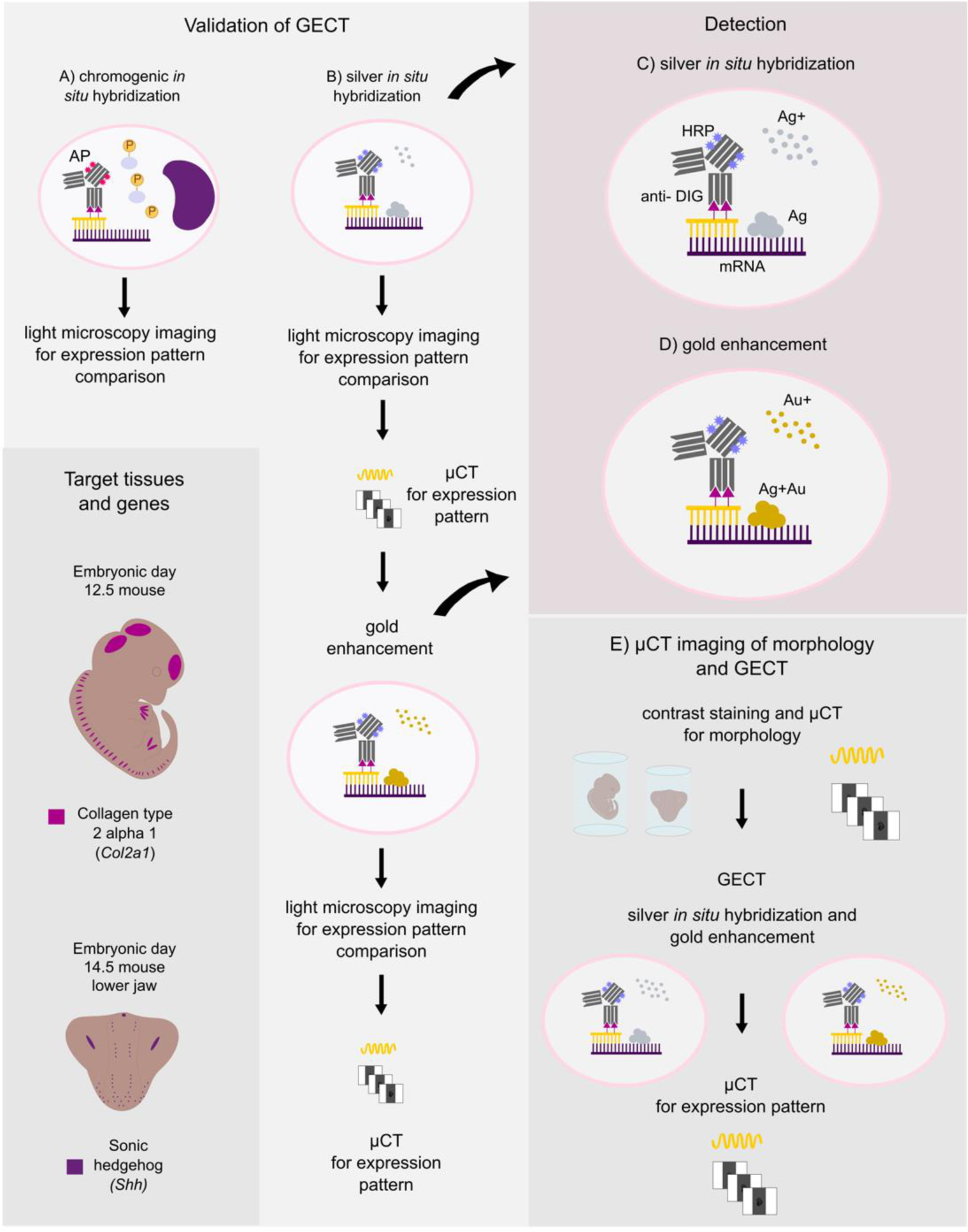
Validation of Gene Expression CT (GECT) and the pipeline for µCT imaging of morphology and gene expression. Validation was conducted by performing chromogenic (A) and silver (B) *in situ* hybridizations in parallel followed by the gold enhancement on the silver *in situ* samples. C) Silver detection was conducted with horseradish peroxidase (HRP) conjugated anti-DIG antibody and standard commercial detection kit (EnzMet, enzymatic metallography, Nanoprobes). During detection, HRP facilitates reduction of silver ions on sites of antibody (and probe) binding. D) Gold enhancement (Gold Enhancement for LM, Nanoprobes) utilizes the silver condensates in the tissue as seeding particles and covers them with elemental gold by reducing gold ions in solution. Resulting signal was imaged with light microscopy and µCT. Some samples underwent µCT after silver *in situ* hybridization (C) and all samples underwent µCT after gold enhancement (D). E) To further validate the detection scheme for routine use, we stained additional samples with phosphotungstic acid, imaged them with µCT to obtain representation of 3D morphology and conducted gene expression detection after µCT imaging for morphology.

## Methods

### Ethics statement

All animal work was performed in accordance with the guidelines of the Finnish Authorization Board under licenses KEK16-021 and KEK19-019.

### Animal work

Wild type NMRI mice (Envigo) were used for collection of tissue samples. Embryonic age was determined as days from the appearance of the vaginal plug (day 0). Embryos were fixed overnight using 4 % paraformaldehyde in 1x phosphate buffered saline (PBS), + 4 °C. Lower jaws were dissected in Dulbecco’s phosphate buffered saline under light microscope before overnight fixation with 4 % PFA in +4 °C. Samples were dehydrated via methanol series and stored in 100 % methanol – 20°C.

### Phosphotungstic acid staining

Samples were rehydrated from 100 % MeOH to 70 % EtOH and stained with 0.3 % phosphotungstic acid (PTA) in 70 % EtOH for 24 – 48 hours in + 8 °C (Metscher, 2009a). After staining, samples were returned to 70 % EtOH. X-ray imaging of PTA -stained samples was conducted in 70 % EtOH using 1.6 mL polypropylene tubes with screw cap and seal (Cryopure, Sarstedt). Samples were stored 70 % EtOH in +4 °C until proceeding with *in situ* hybridization.

### Probe preparation and in vitro transcription

Plasmid construct (Bluescript KS, Strategene) for mouse (*Mus musculus*) collagen type 2 alpha 1 (Col2a1) RNA probes (405 bp) was generated according to Metsäranta et al., 1991, and provided by Mikkola lab, Institute of Biotechnology. RNA probes specific to mouse (*Mus musculus*) sonic hedgehog – gene (*Shh*) exons 1, 2 and 3 were generated using the following primers: CGTAAGTCCTTCACCAGCTTG (forward) and GCTGACCCCTTTAGCCTACA (reverse). cDNA was prepared from mouse embryonic molar tooth RNA (extracted using RNeasy Plus Micro kit, Qiagen) and cDNA constructs were inserted in TOPO II PCR-plasmids using TOPO TA Cloning kit with chemically competent cells according to manufacturer’s protocol (Thermo Fisher Scientific). Prior to *in vitro* RNA synthesis, plasmids were extracted using Miniprep kit (Qiagen).

Plasmids were linearized and probes were prepared as described in Wilkinson & Nieto, 1993 using digoxigenin-conjugated nucleotides (Roche). Specificity of target mRNA binding was confirmed using a sense probe in slide *in situ* hybridization and in whole mount *in situ* hybridization, both chromogenic and metallographic detection were used.

### Whole mount in situ hybridization

Steps were conducted in 24-well plate (Cellstar, Greiner Bio-One) in a volume of 900 µL unless otherwise stated. Samples stored in 100 % methanol were rehydrated to 1x PBS with 0.1% Tween20 (Sigma-Aldrich, P7949-500ML) through a descending methanol series. Ethanol series was used for samples that had been stained with phosphotungstic acid and imaged with µCT prior *in situ* hybridization. After three 15 -minute washes in 1x PBS-T, samples were incubated in 3 % hydrogen peroxide (Honeywell) in 1x PBS-T for 1 h in room temperature (RT). Samples were washed four times with 1x PBS-T for 15 minutes, and then incubated with 10 µg / mL of Proteinase K (Roche, ref 03115879001) in 1x PBS-T, followed by two 10-minute washes with 2 mg / mL glycine (Sigma Life Science, G8790-100G) in 1x PBS-T. After four 15-minute washes with 1x PBS-T, samples were incubated in 4 % PFA in 1x PBS for 20 minutes, followed by three 15-minute washes with 1x PBS-T.

Hybridization step and incubation with antibody were conducted in 2 mL tubes to allow the solution to mix evenly during the incubation period. Hybridization was carried out overnight at + 65 °C with rocking, using probes for mouse *sonic hedgehog* (*Shh*) and *collagen type 2 alpha 1* (*Col2a1*) (Metsäranta et al., 1991) with a concentration of 1 µg / mL. After hybridization, samples were washed two times in 50 % formamide (Emsure, ref 1.09684.1000, Merck), 5x saline sodium citrate (SSC) buffer, 1 % SDS – solution for 30 minutes and 3 times in 50 % formamide, 2x SSC-solution for 30 minutes. All washes were conducted in + 65 °C.

Before antibody incubation, samples were washed in 1x maleic acid buffer (MABT). Anti-DIG antibody conjugated with HRP (Roche, ref 11207733910) was diluted 1 in 50 in solution consisting of 2 % blocking reagent (Roche ref 11096176001), 1 % goat serum (Gibco, ref 16210-064) in 1x MABT. Alkaline phosphatase (AP) conjugated anti-DIG antibody (Roche, ref 11093274910) was diluted 1 in 2000. After addition of the antibody, samples were incubated overnight in + 4 °C with rocking and then washed with 1x MABT for three times 15 minutes, three times 20 minutes, three times 40 minutes and for one hour, RT.

To detect *Shh* and *Col2a1* using alkaline phosphatase activity, BM Purple AP substrate (Roche, ref 11442074001) was added to the samples with AP-conjugated DIG-antibody. Samples were first washed with AP buffer (100mM Tris pH 9.5, 100mM NaCl, 50mM MgCl_2_, 0.1 % Tween20 for 10 minutes and 20 minutes RT before adding the substrate. After substrate addition, samples were incubated and protected from light until strong signal was detected (1-4 hours with *Col2a1* mouse embryos RT, 15-18 hours with *Shh* in mouse embryonic lower jaws, +4 °C). After washing the samples with 1x PBS for 3 x 10 minutes, samples were fixed with 4 % PFA in 1x PBS overnight. Next day samples were transferred to 1x PBS for bright field imaging.

Horseradish peroxidase-based detection was carried out using EnzMet General HRP Detection Kit (Nanoprobes, ref 6010-15mL). Before detection, samples were transferred from 1x MABT to 1:1 solution of 1x MABT and 100 mM sodium citrate buffer, pH 7 for five minutes, and then washed with sodium citrate buffer for 3 x 15 minutes.

Equal volumes of solutions A, B and C of the EnzMet General HRP Detection Kit were added respectively on each sample with anti-DIG antibody conjugated with HRP. 15 -minute incubation was conducted in between each addition of detection solution. After addition of solution C, detection solution was continuously mixed and regularly monitored under light microscope until strong signal was observed and/or first signs of silver precipitate in solution started to emerge (*Col2a1* samples 1-7 minutes, *Shh* samples, 4-10 minutes). After HRP-based detection, samples were transferred to 1 % sodium thiosulphate (Sigma-Aldrich ref 217247-25G) in deionized water and stored overnight in + 4 °C before bright field imaging and gold enhancement. In cases where samples were imaged with µCT before proceeding with gold enhancement (Supplementary Figure 2A), samples were gradually dehydrated to 70 % EtOH after bright field imaging.

### Gold enhancement

Depending on whether the sample was imaged with µCT before gold enhancement, samples were either directly transferred from 1 % sodium thiosulphate to 1x PBS (not µCT imaged, Figures 2-4) or gradually rehydrated from 70 % EtOH to 1x PBS (µCT imaged, Supplementary Figure 2A). Samples were first washed with 1x PBS for 15 minutes and then transferred to 50 mM glycine in 1x PBS for 15 minutes, RT. Before reaction initiation, samples were rinsed in deionized water with 0.1 % Tween20 for 15 minutes, RT. Reaction development was performed following the protocol provided by the manufacturer with small modifications. Equal amounts of solutions A (Enhancer) and B (Activator) of the kit (GoldEnhance for LM, 2112-28ML, Nanoprobes) were mixed and left to incubate for 10 minutes before addition of solutions C (Initiator), D (Buffer) and 0.1% Tween20. Reagent mixture with Tween20 was immediately added on samples, continually mixed, and monitored regularly under microscope until signal was as strong as possible without any precipitate in the solution, varying from 2.5 minutes to 10 minutes. After detection, samples were rinsed in deionized water with 0.1 % Tween20 for five minutes until fixing with 1 % sodium thiosulphate in deionized water with a minimum of 15 minutes to overnight, RT. To test whether gold enhancement could provide additional means to modulate the signal strength in µCT, the procedure was repeated on lower jaw samples after µCT imaging of first gold enhancement signal.

**Figure 2.**
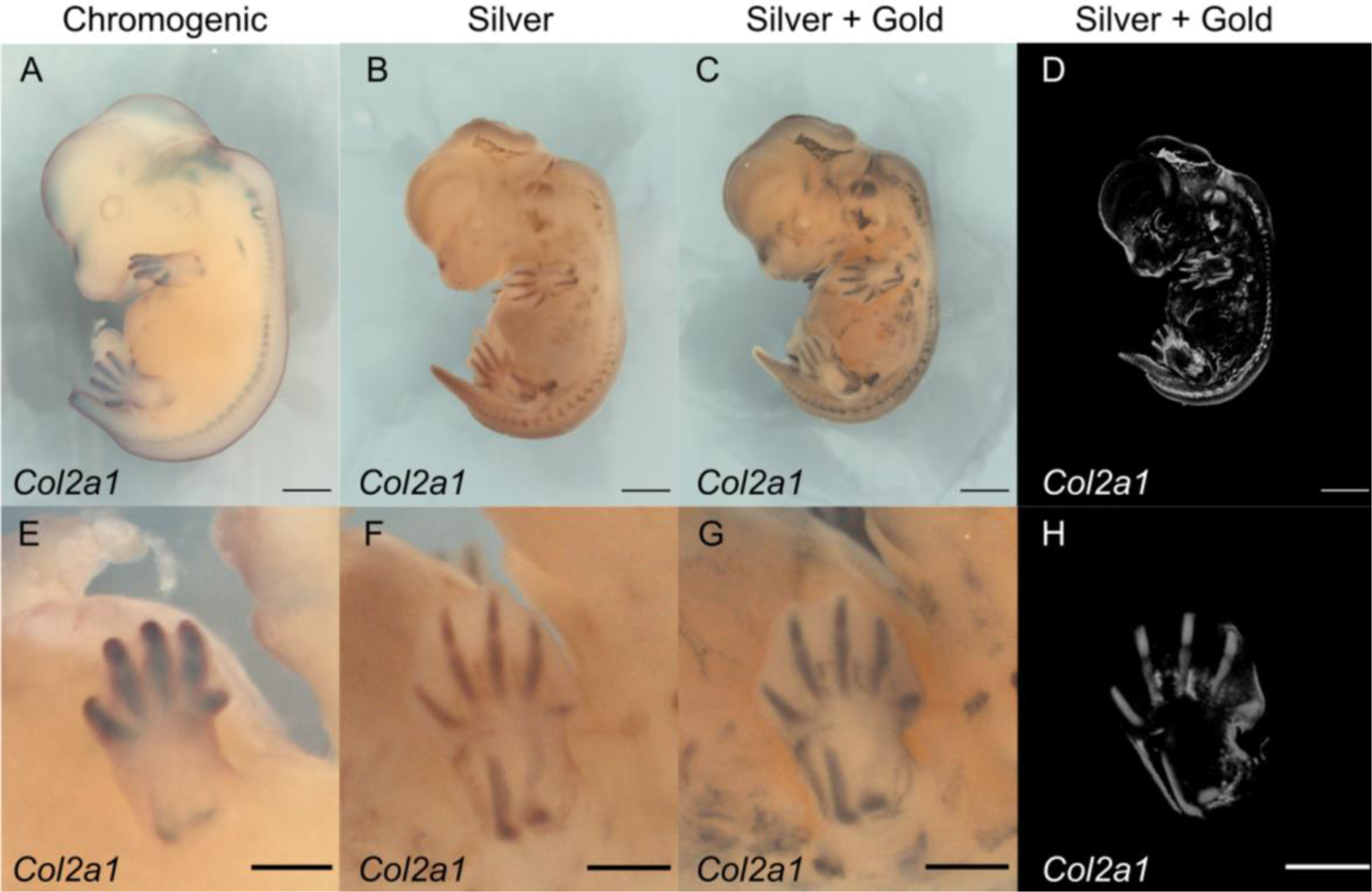
Detection of *Collagen type 2 alpha 1 (Col2a1*) expression with µCT. A) Chromogenic *in situ* hybridization of *Col2a1* in e12.5 mouse embryo. B) Silver *in situ* hybridization of *Col2a1*. C) Gold enhancement of silver *in situ* hybridization shown in B. D) Volume rendering of *Col2a1* expression shown in C. E) Close-up of left upper limb shown in A. F) Close-up of left upper limb shown in B. G) Close-up of left upper limb shown in C. H) Close-up of left upper limb shown in H. Scale 1000 µm in A-D, 500 µm in E-H.

### Vibratome sectioning

To verify that detection components are capable of penetrating to the region of primary enamel knot, jaw specimen undergone silver deposition and gold enhancement was casted in 6 % low melting point agarose (TopVision, ref R0801, Thermo Scientific) in 1xPBS and sectioned to 100 µm sections with vibratome (Microm). Sections were collected in 6-well plate in 1x PBS and imaged over glass slides mounted in aqueous mounting medium (Shandon Immumount, ref 9990402, Thermo Scientific). Vibratome sections were imaged using Zeiss Imager.M2 AX10 with AxioCam HRc camera, EC Plan NEOFLUAR 10x objective and Zeiss Zen2 Pro blue edition software. Images were oriented, cropped, and brightness and contrast were adjusted using ImageJ software (Fiji) (Schindelin et al., 2012).

### Light microscopy imaging

Bright field images of whole mount samples were taken using Zeiss Lumar V12 stereomicroscope, AxiocamICc1 camera, Zeiss Zen 2012 blue edition software and Apolumar S 1.2 x objective with magnification of 25 x and 12 x. Samples were transferred to 1x PBS and set on 4 % low melting point agarose plate to position them. To obtain in focus images of whole sample, multidimensional acquisition (Z-stack) was used with individually optimized focus plane distance and number of focus planes. Extended depth of focus was achieved using highest alignment of “wavelets”-function in Zeiss Zen 2012 blue edition. Images were cropped, and brightness and contrast were adjusted using ImageJ software (Fiji). After bright field imaging, samples were gradually dehydrated to 70 % EtOH for µCT imaging.

### µCT imaging

X-ray imaging was conducted in 70 % EtOH using Bruker Skyscan 1272 µCT device in X-ray laboratory in Department of Physics, University of Helsinki. For tomographic imaging of the overall morphology of whole mouse embryos with a voxel size of 3 µm, we used 0.5 mm aluminum filter, source voltage/current of 70 kV / 142 µA, rotation step of 0.1 degrees and frame averaging of 4. Scanning resulted in 1922 projections over 180° with exposure time of 3038 ms. Total imaging time was approximately nine hours. Samples were stored in 70 % EtOH in + 4 °C until *in situ* hybridization.

µCT imaging after *in situ* hybridization was conducted for lower jaws in 70 % EtOH using the following settings: 1 µm voxel size, no filter, source voltage/current of 30 kV / 212 µA, rotation step 0.1 degrees, frame averaging 5. Scanning resulted in 1877 projections over 180° with exposure time of 1410 ms. Total imaging time was approximately 4.5 hours. For gold enhancement of embryonic lower jaws, a voxel size of 3 µm was used with 0.25 aluminum filter, source voltage/current of 60 kV / 166 µA, rotation step of 0.1 degrees and frame averaging of 5. Scanning resulted in 1922 projections over 180° with exposure time of 1060 ms. Total imaging time was approximately four hours. For µCT imaging of gold enhanced embryos, we used a 3 µm voxel size, 0.25 mm aluminum filter, source voltage/current of 60 kV /166 µA, rotation step of 0.1 degrees and frame averaging of 4. Scanning resulted in 1992 projections over 180° with exposure time of 2269 ms. Total imaging time was approximately seven hours.

### Reconstruction and post-processing

Reconstruction of the µCT data was conducted with nRecon – software (1.7.4.2) with misalignment compensation and ring artifact correction optimized separately for each dataset. Regions of interest were reconstructed as 16-bit tiff stacks that were reoriented and cropped in ImageJ (Fiji) before volume rendering and segmentation in Avizo.

### Segmentation

Segmentation of expression regions was conducted in Avizo (rel 9.0.1) by first excluding 20 % of the darkest pixels in the reconstruction (jaw) and 25 % of the darkest pixels in the reconstruction (whole embryo) by using Thresholding-tool available in Avizo Segmentation-view and then by manually excluding pixels with Brush-tool in Segmentation – view. Pixels were excluded from expression regions based on evaluation where light microscopy data from chromogenic *in situ* hybridization, silver in *situ* hybridization and gold enhancement was considered together with the location of the signal. In other words, high-intensity pixels on the surface of the specimen and in inconsistent locations throughout detections were considered background and therefore excluded. Extracted volume of expression region was derived by assigning a value of 1 for the segmented voxels and a value of 0 for the voxels remaining outside signal region in the reconstruction volume using the Arithmetic–tool in Avizo. Volume rendering and segmentation images were generated with Avizo using Snapshot-tool as 16-bit tiffs. Images were cropped and oriented and brightness and contrast were adjusted in ImageJ (Fiji).

## Results

For the detection of an abundantly expressed gene, we tested silver *in situ* hybridization and gold enhancement for *Col2a1* in embryonic day 12.5 mouse. Chromogenic detection of *Col2a1* (Fig 2A, n=2) is consistent with silver detection and gold enhancement (Fig 2B-C, n=2), showing expression in developing cartilage of limbs, spine and skull. Combination of gold and silver in detection of *Col2a1* enables visualization of expression via µCT (Fig 2D). Expression of *Col2a1* is particularly clear in limbs of the developing embryo, as shown in the upper limb in Figures 2E-H. The silver *in situ* hybridization and gold enhancement generate a clear-cut signal, from which interphalangeal joint spaces are distinguishable as non-expressing tissue (Fig 2F-H).

To test visualization of both the tissue morphology and the gene expression in the same sample, we applied silver *in situ* hybridization and gold enhancement to samples that had undergone phosphotungstic acid (PTA) staining and µCT imaging for morphology. Compared to control samples without PTA staining prior to *in situ* hybridization (Fig 3A-B, n=2), PTA-stained samples replicated the expected expression pattern of *Col2a1* in both stages of detection (Fig 3C-D), similarly with slight increase in background when gold enhancement was applied. µCT reconstruction of the PTA-stained sample prior *in situ* hybridization shows good contrast between the sample and the scanning medium (Fig 3E) and between tissue compartments (Fig 3F). Gold enhanced signal of *Col2a1* expression from the same sample was detectable with µCT (Fig 3G) and segmented expression regions inside the hindlimb are shown in Figure 3H and 3I and Supplementary video 1.

**Figure 3.**
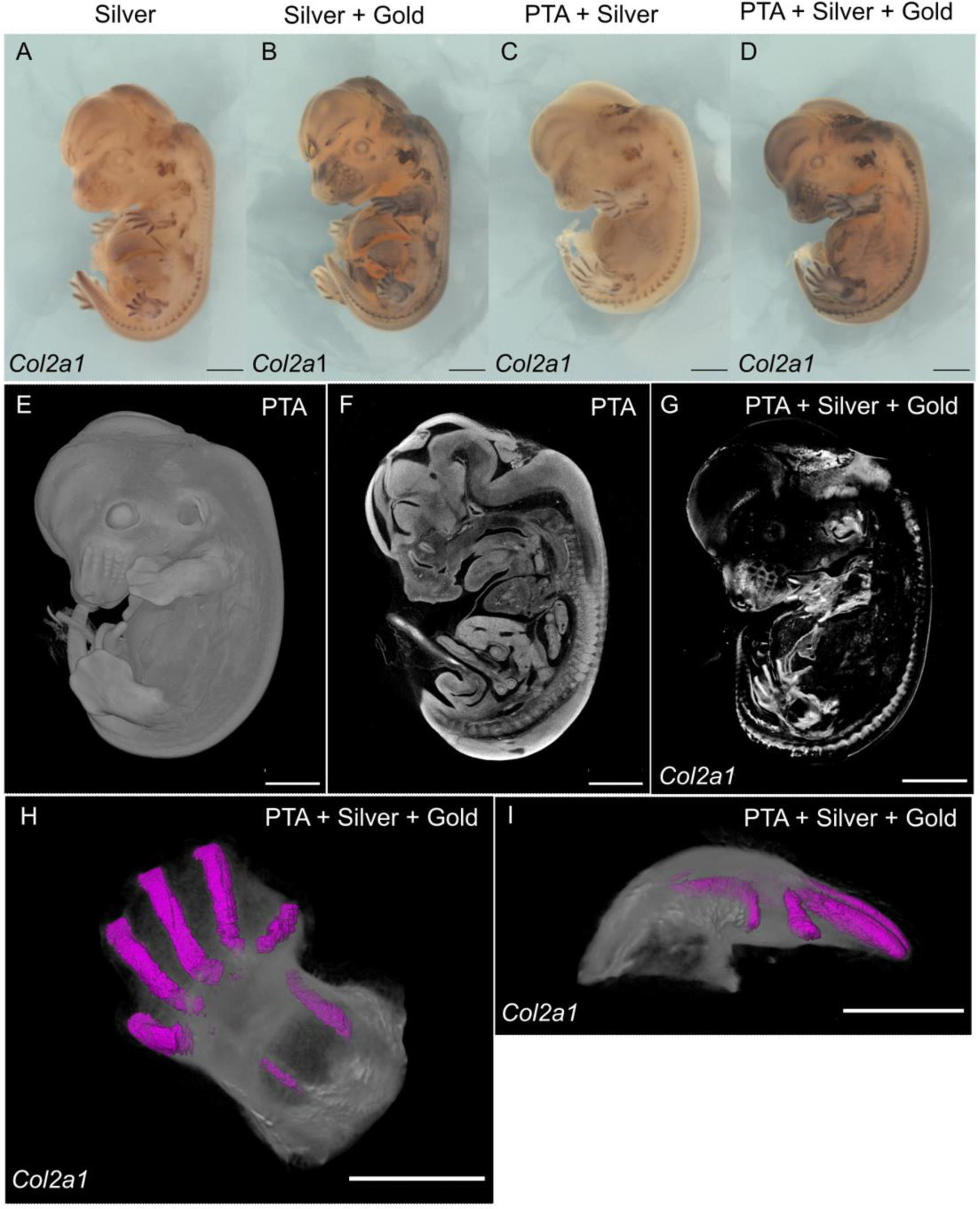
Tissue contrast staining before gene expression detection. A) Silver *in situ* hybridization of *Col2a1* without prior PTA stain. B) Gold enhancement of *Col2a1* detection shown in A. C) Silver *in situ* hybridization of *Col2a1* in mouse embryo stained with PTA. D) Gold enhancement of *Col2a1* detection shown in C. E) Volume rendering of the contrast-stained mouse embryo, prior *in situ* hybridization. F) Cross-section of the mouse embryo shown in E. G) Volume rendering of *Col2a1* expression shown in D. H) Close-up of left hindlimb shown in G. Segmented *Col2a1* expression inside the limb shown in magenta, sagittal view. I) Segmented *Col2a1* expression (magenta) inside left hindlimb, transverse view. Scale 1000 µm in A-G, 500 µm in H-I.

To focus on locally and transiently expressed developmental genes, we examined *Shh* expression in developing mouse lower jaws. In the embryonic day 14 lower jaw samples including tongue*, Shh* expression is detected with chromogenic *in situ* hybridization in enamel knots of the molars, hair and whisker buds of the chin and taste papillae of the tongue. (Fig 4A, n=2). Silver *in situ* hybridization replicates this pattern (Fig 4B, n=3) and gold enhancement of the silver detection preserves it (Fig 4C, n=3). Silver *in situ* hybridization performed equally to chromogenic detection in every trial (see Supplementary Table 1). We also observed that *Shh* expression regions detected with silver *in situ* hybridization and gold enhancement are more spatially defined compared to the signal produced by chromogenic detection (Fig 4A-C, see also Supplementary Figure 1).

**Fig 4.**
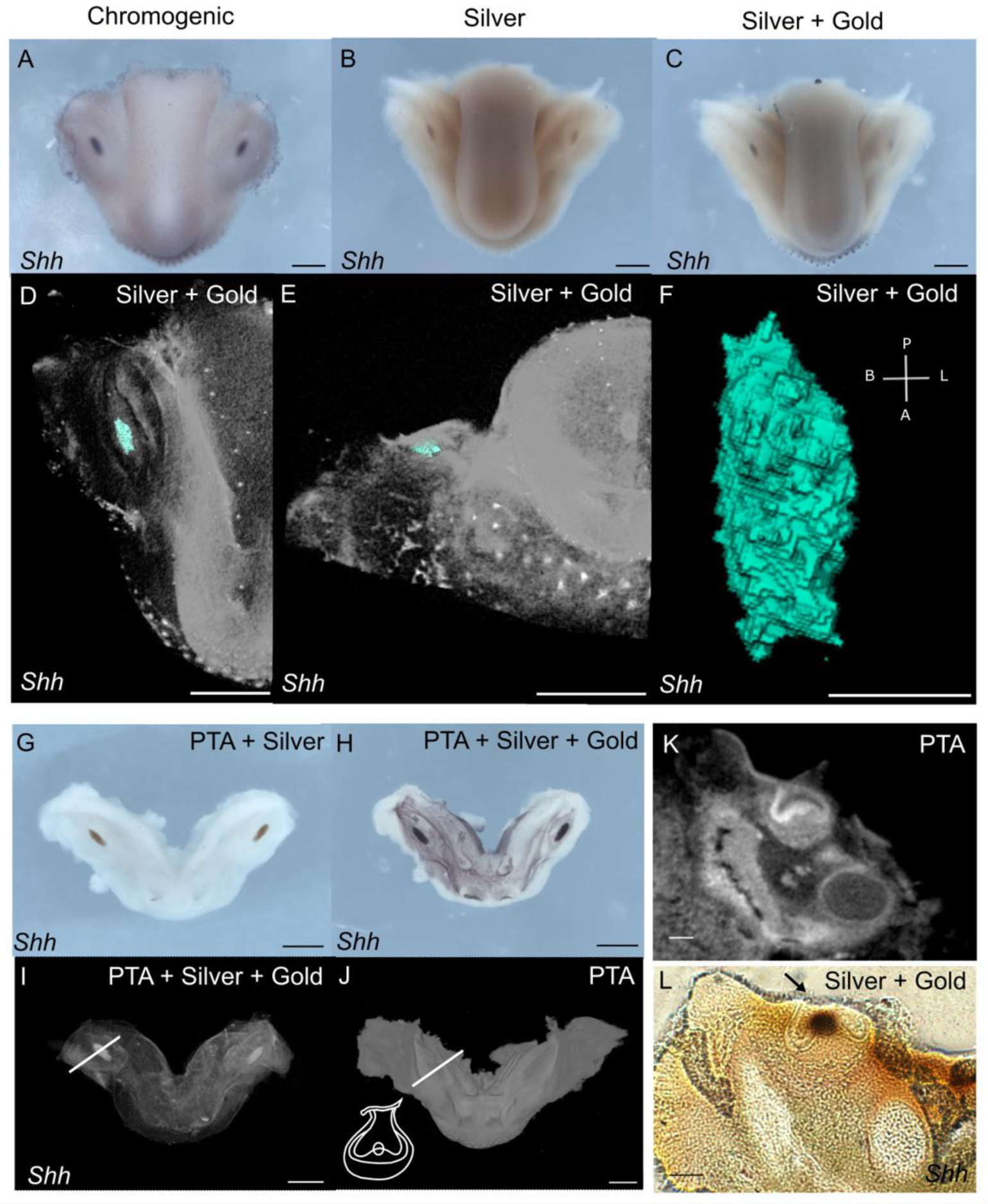
µCT visualization of *Sonic hedgehog (Shh)* expression from unstained and contrast-stained tissues. A) Chromogenic *in situ* hybridization of *Shh* expression in mouse e14.5 jaw. B) Silver *in situ* hybridization of *Shh*. C) Gold enhancement of *Shh* detection shown in B. D) Volume rendering of *Shh* expression from µCT data, occlusal view. Expression in embryonic molar shown in teal. E) Volume rendering of *Shh* expression, expression in embryonic molar shown in teal. frontal view. F) Segmented expression region of *Shh* in mouse molar, occlusal view. B = buccal, L = lingual A = anterior, P = posterior. G) Silver *in situ* hybridization for *Shh* applied to contrast-stained jaw. H) Gold enhancement of *Shh* detection shown in G. I) Volume rendering of µCT data from jaw shown in H. J) Volume rendering of contrast stained jaw shown in G, H and I, prior Shh detection. White line shows approximate section plane shown in K. Structures of a developing molar in shown plane are visualized in schematic in the lower left corner. K) Frontal virtual section of developing molar stained with PTA. EK = enamel knot. L) Frontal vibratome section of jaw undergone silver *in situ* hybridization and gold enhancement for *Shh* (dark brown). EK = enamel knot. Epithelial tearing shown with arrow. Scale 500 µm in A-E, 100 µm in F, 500 µm in G-J, 100 µm in K-L.

µCT imaging of jaw samples was conducted after both silver *in situ* hybridization and gold enhancement. µCT imaging of silver detection resulted in no observable attenuation on sites of *Shh* expression and imaging after first gold enhancement round visualized expression regions only partially. (Supplementary Figure 2). In samples containing tongue, all expression sites were visible with µCT after repeating gold enhancement (Fig 4D-F, *Shh* in enamel knot shown in teal color).

Similarly to the *Col2a1*, *Shh* expression was detectable in the tissues stained with PTA (Fig 4G-H, n=2) and visible in µCT (4I, n=2). We note that whereas PTA staining before *in situ* hybridization provided contrast for µCT detection of jaw tissue (Fig 4J, 4K), PTA is washed away from tissues during *in situ* hybridization (Fig 4I).

*In situ* hybridization treatments cause tissue distortion, which may affect 3D reconstructions. When compared to frontal virtual section of developing molar from jaw stained with PTA and imaged with µCT (Fig 4K), frontal vibratome section of a whole mount sample undergone *in situ* hybridization and gold enhancement shows tearing of surface epithelium on site of dental chord (arrow, Fig 4L). Expression region of *Shh* in whole mount samples is consistent with section *in situ* hybridization (Supplementary Fig 1). Epithelial tearing occurred frequently in jaw samples and also in samples undergone chromogenic detection, and is thus not specific to the steps of gold enhancement. We observed no drastic tearing or distortion of tissues in whole embryo samples on sites of expression. Control sample without prior PTA staining shown in Fig 3A and B exhibits the extent of tearing in stomach region that occurred during *Col2a1* detection. Samples stained with PTA prior detection show slightly less tissue tearing (Fig 3C-D).

## Discussion

Despite the wide range of resolution available in µCT imaging, there remains the challenge to image specific molecules or gene activity. Initially peroxidase-assisted reduction of silver was used to visualize distribution of collagen type II and acetylated alpha-tubulin proteins in chick embryos with silver (Metscher & Müller, 2011). This approach utilized conventional immunohistochemical detection of proteins, in which antibody is conjugated with horseradish peroxidase (Powell et al., 2007). Here we apply a similar, straightforward approach to detection and visualization of mRNA distribution in whole mount samples.

In addition to the lack of molecular markers, contrast agent use and development for µCT imaging is still regarded to be in its early stages compared to other imaging modalities (Rawson et al., 2020). Here our aim was to develop a method that can be incorporated into existing pipelines of i) µCT imaging of soft tissue ii) detection of gene expression with *in situ* hybridization. We show that the commonly used phosphotungstic acid staining can be performed prior to GECT. With two rounds of µCT imaging it will be possible to obtain first the tissue structure and then, after the *in situ* hybridization, gene expression pattern in 3D.

*In situ* hybridization is a robust, widely used method for spatial detection of gene expression and there exists a variety of optimized protocols for a range of model organisms and both for tissue sections and whole mount samples (Acloque et al., 2008; Hafen et al., 1983; Hemmati-Brivanlou et al., 1990; Rosen & Beddington, 1993; Schulte-Merker et al., 1992; Thisse & Thisse, 2008; Yu & Holland, 2009). The pipeline presented here requires relatively small changes to the existing *in situ* hybridization pipelines. Compared to the commonly used chromogenic detection of gene expression patterns, silver *in situ* hybridization provides more precise spatial readout of gene expression (Fig 2 and 4, Supplementary Figure 1, (Powell et al., 2007). An added advantage is that if µCT imaging is not required, the pipeline can be stopped after the silver detection and the samples can be sectioned for histological analyses.

Areas benefitting from further developments include tissue distortion caused by *in situ* hybridization. Whereas in developing mouse molars there is tearing of epithelium after *in situ* hybridization, whole mouse embryos hold their shape better, exhibiting only a slight tearing in the stomach. It is conceivable that these are not caused by any distinct step in the protocol but stem from repeated processing of the samples. One easily modifiable aspect would be the length and number of dehydration and rehydration steps preceding µCT and *in situ* hybridization. Time required for sample collection, tissue staining, µCT imaging of morphology, *in situ* hybridization and µCT imaging of gene expression presented in this proof-of-concept work is approximately two weeks and it has room for further developments. Based on different responses of the samples we used, we additionally suggest that distortion is somewhat dependent on sample size and type. Specific knowledge of the target tissue allows further optimization of the detection conditions.

We observed modest superficial background signal in the samples with silver detection, consistent with the observations of Metscher & Müller (2011). This background became stronger alongside the expression signal when the same samples were processed through gold enhancement (Fig 2, Fig 3, Fig 4). The benefit of µCT imaging in this context is also that the signal visible on the surface of the sample can be discriminated from the signal within the sample. Nevertheless, we recommend that at least when working with unfamiliar probes and genes known to have superficial expression, one should run chromogenic detection alongside the combination of silver and gold detection to be able to distinguish the background from the signal.

Metscher and Müller propose a quantification method for the level of gene expression based on the intensity values of µCT imaging data (Metscher & Müller, 2011). We note that silver *in situ* hybridization is an enzyme-based detection method and inherently includes variation in the intensity of the signal not necessarily reflecting the biological state of the tissue. Gold enhancement is not based on enzymatic activity and seems to replicate the signal of silver detection faithfully, and it remains to be explored how well the method captures differences in gene expression levels.

In conclusion, the protocol developed in this study should help to integrate µCT imaging and detection of gene expression using in situ hybridization, two commonly used methods that when combined, should provide new opportunities for biological imaging.

## Acknowledgements

We thank J. Ollonen and members of the Center of Excellence in Experimental and Computational Developmental Biology for advice and discussions. We thank O. Hallikas, T. Rajala, K. Väänänen and L. Milocco for commenting the manuscript. We thank A. Viherä, Merja Mäkinen and Raija Savolainen for technical assistance.

